# NMR structure of a vestigial nuclease provides insight into the evolution of functional transitions in viral dsDNA packaging motors

**DOI:** 10.1101/2020.07.06.188573

**Authors:** Bryon P. Mahler, Pawel J. Bujalowski, Huzhang Mao, Erik A. Dill, Paul J. Jardine, Kay H. Choi, Marc C. Morais

## Abstract

Double-stranded DNA viruses use ATP-powered molecular motors to package their genomes. To do so, these motors must efficiently transition between initiation, translocation, and termination modes. Here, we report structural and biophysical analyses of the C-terminal domain of the bacteriophage phi29 ATPase (CTD) that suggest a structural basis for these functional transitions. Sedimentation experiments show that the inter-domain linker in the full-length protein promotes dimerization and thus may play a role in assembly of the functional motor. The NMR solution structure of the CTD indicates it is a vestigial nuclease domain that likely evolved from conserved nuclease domains in phage terminases. Despite the loss of nuclease activity, fluorescence binding assays confirm the CTD retains its DNA binding capabilities and fitting the CTD into cryoEM density of the phi29 motor shows that the CTD directly binds DNA. However, the interacting residues differ from those identified by NMR titration in solution, suggesting that packaging motors undergo conformational changes to transition between initiation, translocation, and termination.

## INTRODUCTION

A fundamental step in the life cycle of any virus is encapsidation of the viral genome within a protein shell, or capsid, which protects the genome from environmental assault as the virus transits between hosts. There are two basic strategies that viruses use for genome packaging. In one strategy, the genome first condenses, typically via interaction with virally encoded positively charged proteins. The virus capsid then assembles around the condensed genome. This strategy is used by many enveloped RNA viruses such as alpha viruses and flaviruses. In a second strategy, an empty virus shell is first assembled, and the genome is then actively packaged into this pre-formed container. This is the strategy used by some ssRNA (*1*) and ssDNA viruses (*2*, *3*), and virtually all dsDNA viruses such as herpes virus, pox virus, adenovirus, and all the tailed dsDNA bacteriophages (*4*). This second strategy is remarkable considering the enthalpic, entropic, and DNA bending energies that must be overcome to package DNA to near crystalline densities within the confined space of the capsid.

The molecular motors that power genome encapsidation are some of the most powerful molecular motors in nature, capable of producing forces in excess of 50 piconewtons (*5*). To provide a frame of reference, approximately 5 pN are necessary to break a hydrogen bond, and approximately 200 pN are required to break a weak covalent bond. For comparison, myosin and kinesin each operate at around 5 to 10 pN (*6*, *7*). Thus, viral dsDNA packaging motors operate at high forces, as they must to overcome the ~20 atm of pressure estimated to be present within fully packaged capsids. The energy for packaging is provided by a virus-encoded ATPase that converts the chemical energy of ATP hydrolysis into the mechanical translocation of DNA. These ATPases belong to a large ancient family of ring ATPases involved in various polymer movement/manipulation tasks such as cell division, protein degradation, chromosome segregation, DNA recombination, strand separation, and conjugation. Hence, the operation of viral DNA packaging ATPases may also reflect the mechanistic principles of a broad class of molecular motors responsible for basic macromolecular partitioning processes.

Further, successful DNA packaging requires that motors carry out several related yet functionally distinct tasks. *During initiation*, the packaging motor must self-assemble at a unique vertex of the capsid and recognize its own DNA amongst the multitude of other DNAs present in the cell. *During translocation*, the motor must coordinate the activities of multiple components to efficiently translocate DNA. *During termination*, the motor transitions from a translocation-mode to a termination-mode, wherein the highly pressurized DNA is retained in the head, and the motor disengages to initiate packaging on another empty procapsid. Thus, dsDNA viruses must execute a complex sequence of carefully orchestrated molecular tasks to successfully encapsidate their genomes. While each of these tasks have been examined individually to varying extents, very little is known regarding how dsDNA packaging motors transition between them.

Bacteriophage phi29 has long served as a model system for investigating genome packaging, and a highly efficient in vitro phi29 DNA packaging system has been developed that has allowed packaging to be interrogated via multiple experimental approaches (*4*, *8*, *9*). Genetic, biochemical, and structural studies indicate that the motor is comprised of three macromolecular components (Fig 1A) (*4*, *10*): 1) a dodecameric portal, or connector protein (gene product 10 (gp10)) (*11*); 2) a pentameric ring of a phage encoded structural RNA molecule (pRNA) (*11*–*15*); and 3) a pentameric P-loop ASCE ATPase ring (gene product 16 (gp16)), analogous to the large terminases in other phage systems, and which provides the energy for packaging (*10*, *11*, *16*). These three components are arranged as three co-axial rings, and the dsDNA genome is translocated into the phage capsid through a continuous channel along their common central axis (Fig 1A). Note that unlike other bacteriophages that produce a concatemeric genome, polymerases from phi29-like phages use a protein-priming mechanism to produce a unit-length genome capped at both 5’-ends (*17*). Hence, phi29-like phages would not seem to need the additional nucleolytic machinery that is present in other phages, and which is used to cut the concatenated genome at the beginning and end of packaging.

**Figure 1.**
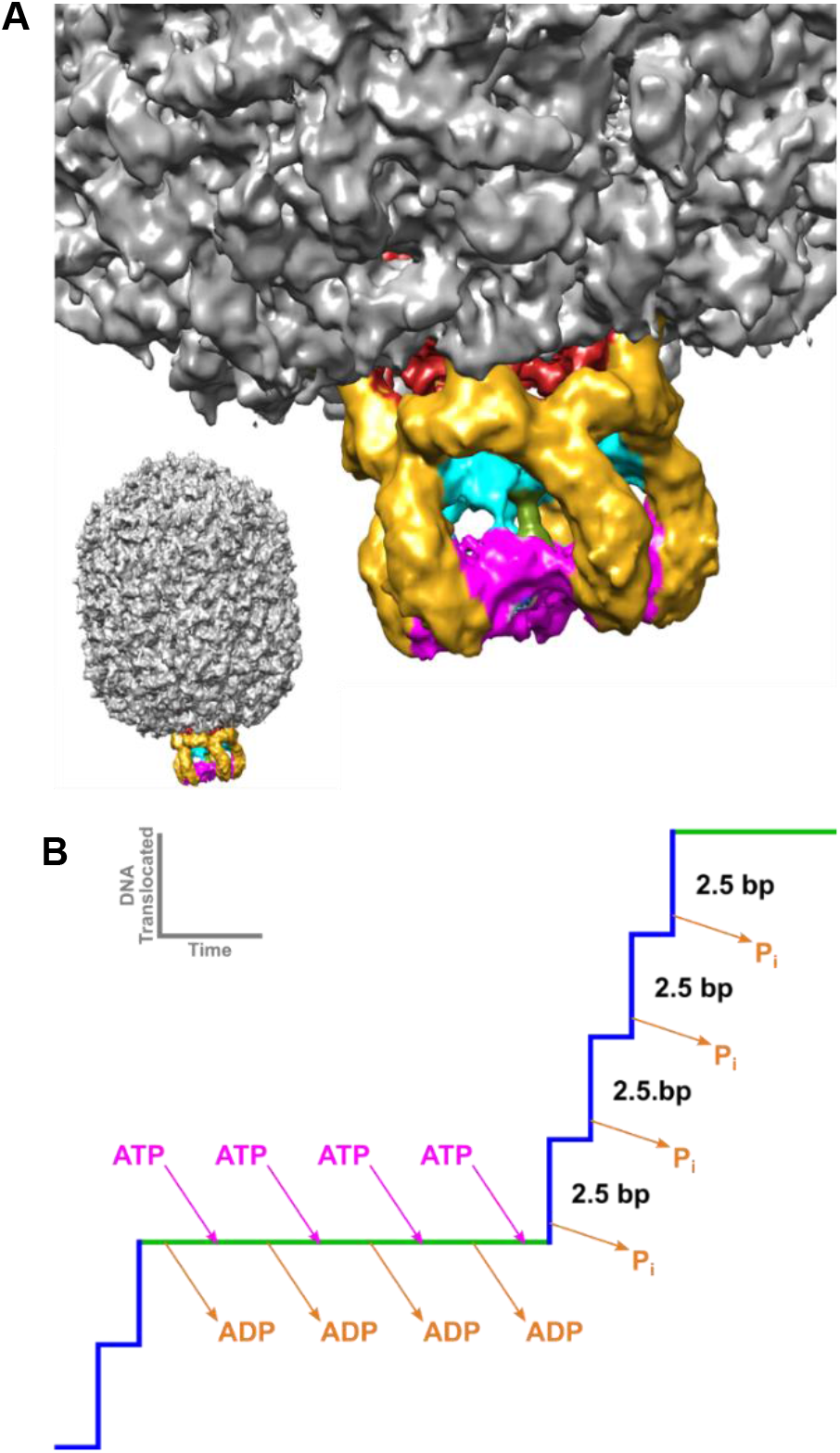
Packaging Motor of the Bacteriophage phi29. *(A) A cryoEM reconstruction of the phi29 ATPase motor complex attached to the procapsid* (24)(EMDB: EMD-6560). The connector, pRNA, CTD, NTD and dsDNA are colored red, yellow, cyan, magenta, and green, respectively. (B) Schematic model of a single dwell-burst packaging event in the mechanochemical cycle of the phi29 packaging motor. During the dwell phase (green line), the ATPase motor is charged with ATP. During the subsequent burst phase (blue line) hydrolysis of the ATP results in the translocation of 10 bp of dsDNA in four 2.5 bp sub-steps associated with 4 hydrolysis events (21).

Biochemical and single molecule analysis indicate that the phi29 genome packaging motor operates in a complex, highly coordinated fashion during translocation (*18*–*21*), and that the mechano-chemical cycle is separated into a dwell phase and a burst phase (Fig 1B) (*21*). No translocation occurs during the dwell, as all five gp16 subunits release ADP from the previous hydrolysis cycle and load ATP in an interlaced manner. More recent laser tweezer experiments show that two distinct regulatory mechanisms coordinate nucleotide exchange during the dwell (*22*). During the subsequent burst, four gp16 subunits sequentially hydrolyze ATP, resulting in translocation of 10 bp of DNA in four 2.5 bp sub-steps. The role of the fifth subunit is not entirely clear, but it has been proposed to play a functionally unique regulatory role in aligning the motor with DNA for the next translocation burst (*18*, *20*). Using altered DNA substrates, additional single molecule experiments suggest that the motor makes two distinct types of contact with the translocating DNA (*23*); during the dwell phase, the motor makes specific electrostatic contacts with the phosphate backbone of the DNA, whereas the motor uses non-specific contacts during the burst to actively translocate the DNA. It has also been shown that DNA rotates 14° during each 10 bp translocation burst, likely to maintain motor/substrate alignment, and that the magnitude of this rotation is coupled to the changing step-size of the motor as the head fills (*20*).

While extensive biochemical and single molecule analysis have provided a detailed kinetic scheme describing what happens during packaging (Fig 1B), the molecular motions that underlie force generation and subunit coordination during translocation remain largely unknown. Additionally, how the motor transitions between translocation, termination, and initiation mode is similarly poorly understood. To visualize these events, we have thus been pursuing a hybrid structural approach wherein individual motor components are solved to atomic resolution via X-ray crystallography and then fitted into more moderate resolution cryoEM reconstructions of the entire complex (*12*, *24*). We have previously determined atomic resolution structures of every component of the motor except for the ~120 amino-acid C-terminal domain (CTD) of the packaging ATPase gp16 (*11*, *13*, *24*). Here, we describe the atomic structure of the CTD as determined by solution NMR. This structure, along with atomic structures of the connector, the pRNA, and the NTD of gp16, was fitted into a cryoEM reconstruction of phi29 particles stalled during packaging (**doi:** https://doi.org/10.1101/2020.05.23.112524), resulting in the first complete near-atomic resolution model of an actively packaging motor. Additionally, analytical ultracentrifugation, fluorescence-based binding assays, and NMR chemical shift perturbation (CSP) experiments were used to characterize ATPase assembly and nucleic acid binding by the CTD. Together, these results show how the CTD interacts with other motor components and the translocating DNA and provides insight into how viral dsDNA packaging motors transition between genome processing and translocation functions.

## RESULTS

### Solubility and oligomerization of the CTD of gp16

The phi29 DNA packaging ATPase is a 39-kDa 332-amino acid protein consisting of two domains separated by a long linker (*4*). The N-terminal domain is approximately 200 amino acids long and corresponds the conserved ASCE ATPase domain. The last ~ 120 amino acids were predicted to adopt an oligonucleotide/oligosaccharide binding fold (OB-fold), implying a possible role in either pRNA or DNA binding (*4*). Attempts to crystallize full-length protein have thus far been unsuccessful. Hence, to potentially crystallize the two domains separately, gp16 was subjected to limited trypsinization (*24*). Briefly, we found that trypsin cut only once in the inter-domain linker region, producing two fragments, one corresponding to the N-terminal ASCE ATPase domain (residues 1-207) and the other corresponding to the inter-domain linker and the CTD (residues 208-332).

While the N-terminal domain grew crystals that yielded an atomic resolution structure (*24*), exhaustive attempts to crystallize the construct corresponding to the CTD and the interdomain linker were unsuccessful. However, the dispersion of peaks in an HSQC spectrum of the construct indicated that the protein was well-folded, stable during NMR data collection, and thus a good candidate for structure determination by solution NMR (Fig S1A). Unfortunately, the construct was incapable of consistently reaching concentrations of 250 μM, necessitating long NMR data collection times that nonetheless resulted in sub-optimal signal intensity. Additionally, chemical shifts in the N-terminal tail (corresponding to the interdomain linker in the full-length protein) were present as multiple peaks, readily apparent in G211 and G215 (Fig S1B), indicative of slow exchange on the NMR time scale or multiple species present in the sample. The intensity of minor peaks increased over time with a corresponding decrease in the major peak, suggesting that the multiple peaks resulted from proteolytic cleavage and thus represents multiple species in the sample (Fig S1C). The heteronuclear NOE (hnNOE) data, which is sensitive to fast dynamic motions, showed that the first 20 residues are very flexible (Fig S2A), suggesting that these residues are susceptible to proteolysis. Fortunately, the flexibility of the tail residues also suggested that they were not part of the core structure and thus could be removed without affecting the global fold.

Thus, to further optimize the sample for NMR structure determination, four additional clones were made where the first 5, 10, 15, or 20 amino acids were removed. The clone where the first 20 amino acids were removed did not express well in *E. coli.* In contrast, the construct with 15 residues removed, including residues 223-332, produced good yields and was thus chosen for structure determination. Since this construct was the shortest construct that could be expressed, we will refer to it herein as the CTD construct to differentiate it from the longer construct that included the interdomain linker region (CTD-L). The HSQC of the CTD also indicated a well folded protein and lacked the minor peaks present in longer CTD-L (Fig 2). The first 4 residues of the CTD were still flexible, with hnNOE values less than 0.65 (Fig S3A). The 15 residues that are removed in this construct are very acidic, resulting in an increase in the theoretical pI from 6.2 to 8.0 after removal of these residues. This change in pI made possible the use of more acidic buffers while still maintaining a pH far from the pI. Thus, to further optimize NMR data collection conditions, we explored the stability and solubility of the protein in a number of different buffers using protein thermal shift (PTS) assays (data not shown). Remarkably, switching to a pH 6.0 MES buffer allowed us to able obtain concentrations up to 1 mM for this construct, further facilitating NMR structure determination.

**Figure 2.**
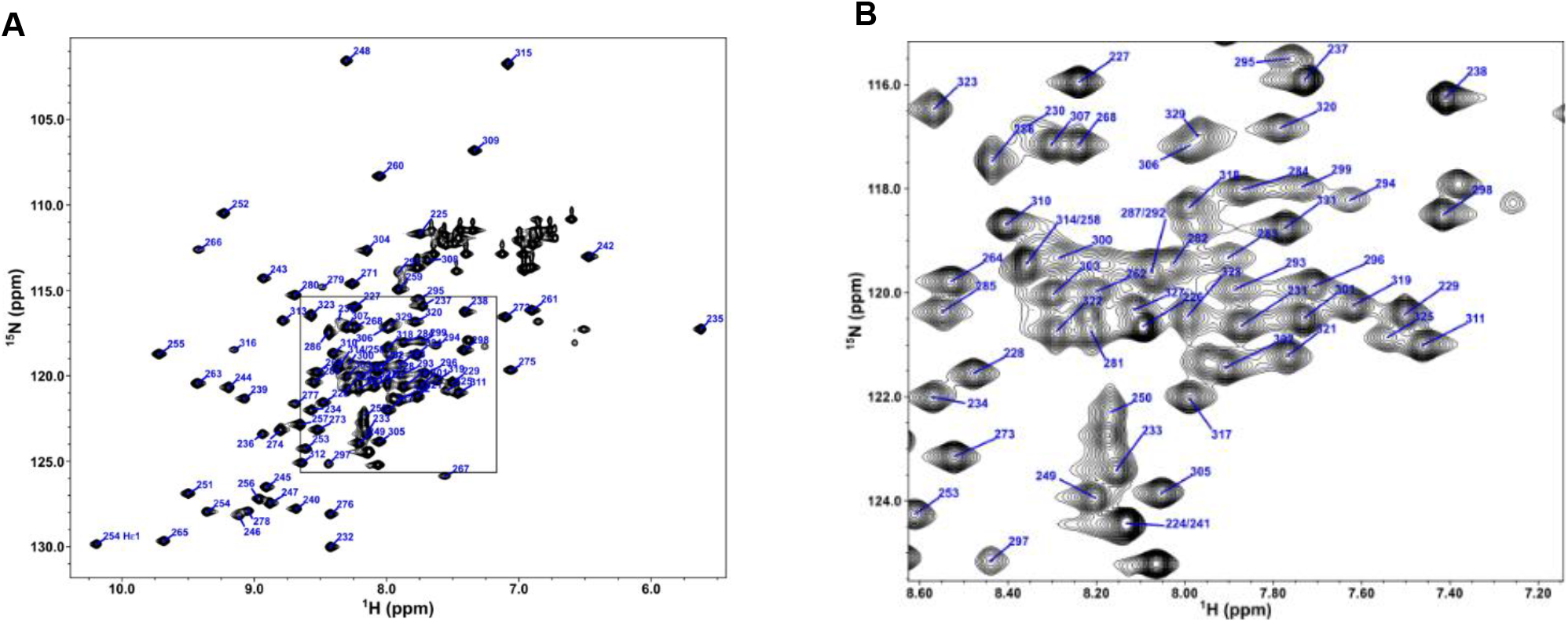
NMR Spectrum of phi29 gp16 (223-332). (A) The 1H-15N HSQC of the phi29 gp16 (223-332) with backbone peaks labeled by residue number. The tryptophan side chain epsilon proton-nitrogen pair is also labeled. (B) Inset of the HSQC from the boxed region in (A)

Longer than expected rotational correlation times for CTD-L suggested that dimers of the construct might be present in solution. At 250 μM, CTD-L has a rotational correlation time of 13 ns, which is much higher than expected for a 14 kDa protein. Conversely, the shorter CTD had a correlation time that was more consistent with a monomer at 9 ns for 500 μM of protein. Thus, to characterize the oligomeric states of the constructs, we examined their sedimentation velocities via analytical ultracentrifugation (SV). SV results for CTD-L showed the percentage of dimers in solution increases from ~15% to ~25% at monomer concentrations of 25 and 55 μM, respectively (Fig 3A, C). In contrast, we observed a constant population of only 2% dimers for the shorter CTD at concentrations of 30 and 85 μM (Fig 3B, C). These results indicate that the N-terminal tail of the CTD-L (the inter-domain linker in the full length gp16) induces dimerization. Since the full-length packaging ATPase in phi29 forms a pentamer when assembled on the procapsid, these results suggest that the linker between the N- and C-terminal domains plays a role in oligomerization of the functional complex.

**Figure 3.**
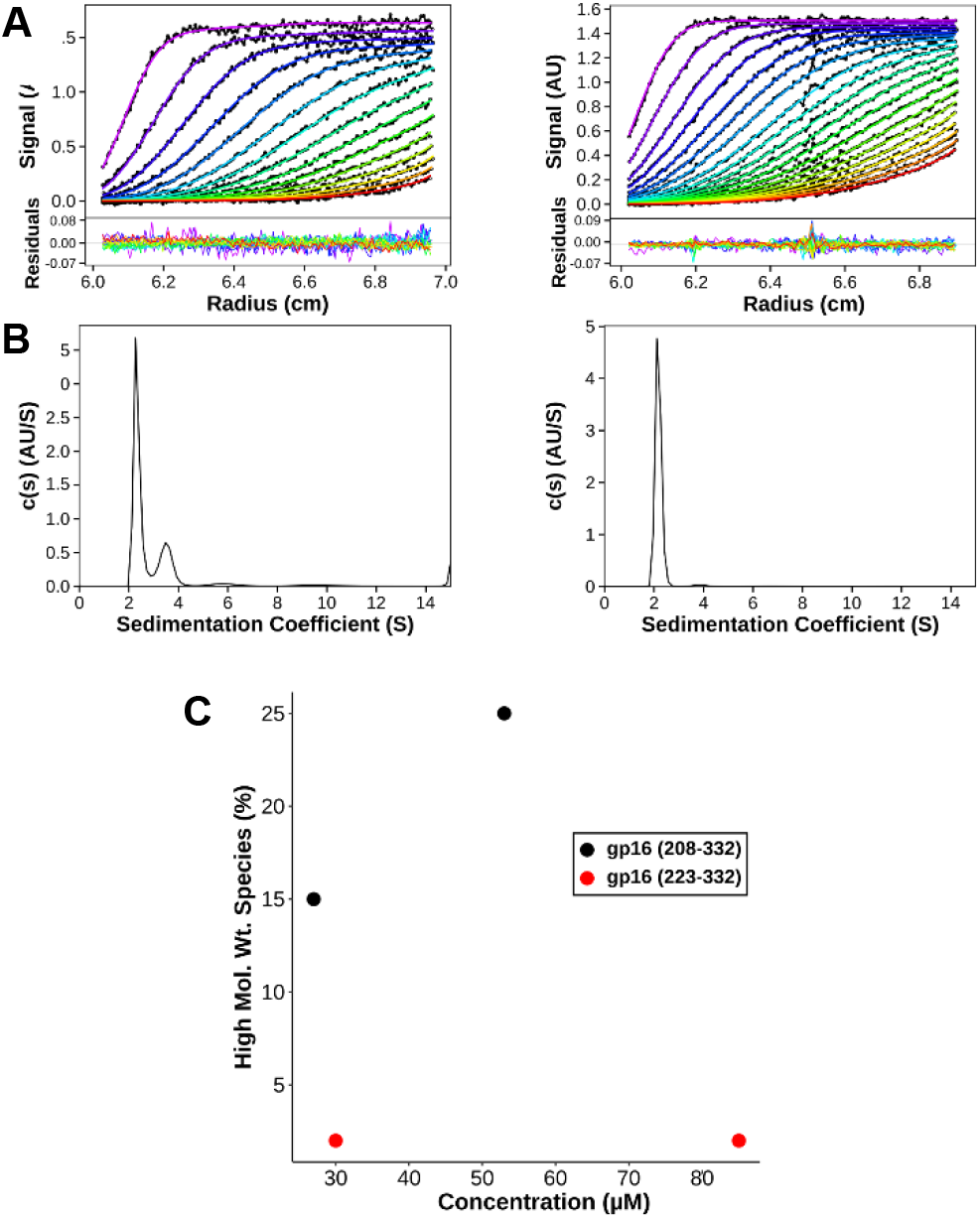
Oligomerization of CTD. (A) SV data and continuous sedimentation distribution of gp16 (208-332) at a concentration 53 μM. (B) SV data and continuous sedimentation distribution of gp16 (223-332) at a concentration of 85 μM. (C) Plot of percentage of a higher molecular weight species with respect at various protein concentrations. The higher molecular weight species is defined as the area under the peak with a sedimentation coefficient around 3.5 for gp16 (208-332) and around 3.8 for gp16 (223-332).

### NMR solution structure of the gp16 CTD

A nearly complete resonance assignment (93%) of the non-proline backbone amides was obtained for the CTD (Fig 2). The longest contiguous stretch of missing assignments was from M288 to I290, likely due to motion on the intermediate time scale in the flexible loop spanning residues D282-N296 as indicated by the high spin-spin, or R_2_, relaxation rates in the region (Fig S3B). Simulated annealing with 776 experimental restraints, including 578 NOE distance restraints, produced a 15-structure ensemble with a secondary structure backbone RMSD of 0.58 Å (Table S1, Fig S3C). The resulting structure is a small compact α/β-fold measuring approximately 34 by 40 by 27 Å and consisting of a 4-stranded β-sheet flanked on one side by two α-helices and two stretches of coil on the opposite side (Fig 4A; PDB 6V1W). One coil connects the N-terminus of the construct to the first β-strand and corresponds to the C-terminal end of the NTD-CTD linker in the full-length protein; the other coil connects two strands on opposite sides of the central β-sheet. The first three strands of the domain are connected by short β-hairpins, and thus adopt an anti-parallel arrangement. The fourth strand (β4) is parallel to the first (β1) and is connected to the third (β3) by the loop that crosses back over strands β1 and β2. The first α-helix (α1) is connected to the end of β4 by a long flexible loop, and to α2 by another loop that runs approximately parallel to strand β3 (Fig 4B). Calculating the electrostatic surface potential indicates two positively charged patches on opposite corners of the central sheet, consistent with a possible role in binding either the pRNA and/or the DNA during DNA packaging (Fig 4C). Coloring residues on the molecular surface according to the Kyte-Doolittle hydrophobicity scale indicates that there is a deep cavity with a hydrophilic rim formed by the two helices and a hydrophobic floor corresponding to the face of the β-sheet flanked by the two helices (Fig 4D).

**Figure 4.**
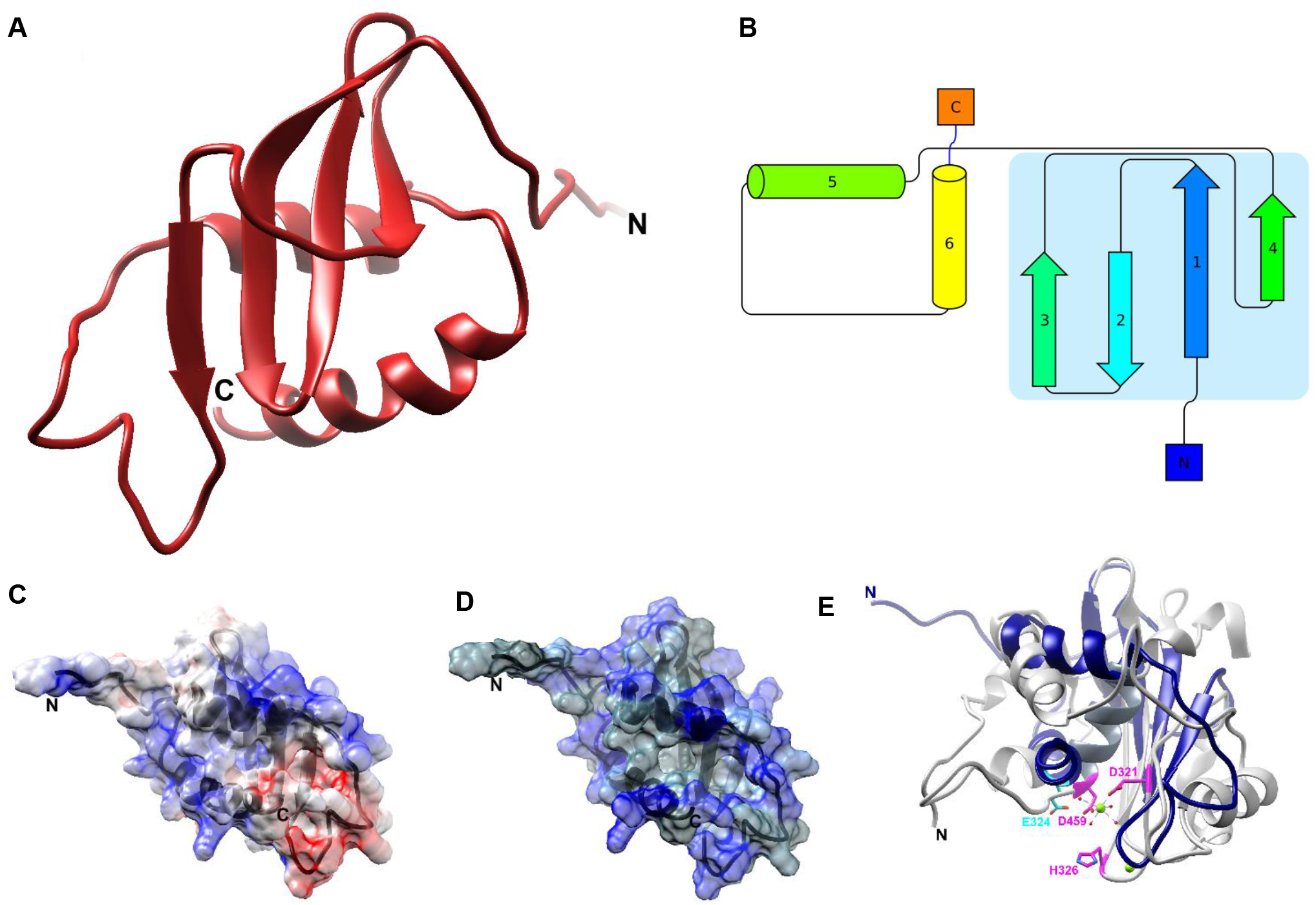
CTD Structure and Surface Properties. (A) Ribbon diagram of the lowest energy structure in the ensemble. (B) Topology diagram of the CTD, with the central β-sheet outlined in light blue. (C) Electrostatic surface potential of the CTD. Red represents negatively charged patches and blue positively charged patches. (D) Hydrophobic surface map of the CTD. Blue represents polar residues, while gray represents hydrophobic regions. (E) Superposition of the CTD (blue) with the C-terminal nuclease domain from bacteriophage P22 (grey; PDB 4DKW). Catalytic residues in P22 are colored pink. There is little apparent sequence homology of the catalytic/metal binding nuclease residues in the CTD; E324 from the CTD (shown in cyan) is the only acidic residue near the P22 magnesium ion (shown in green), but does not correspond to any of the conserved catalytic/metal binding residues in the nuclease domain.

### Identifying homologs of the CTD of gp16

A BLAST search of the CTD sequence does not identify any obvious homologues other than packaging ATPase CTDs for close relatives of phi29. However, since structural homology often persists over long evolutionary distances where sequence homology vanishes, we used the NMR structure of the CTD to search for proteins/domains with similar structures. The Dali server (*25*) returned several structural homologs of the CTD, all belonging to the RNase H superfamily. The common core of the RNase H fold is a 5-stranded β-sheet sandwiched between α-helices on either side of the sheet. While variations on this basic architecture are present in different branches of the superfamily, the most common topology is a mixed β-sheet consisting of 5 β-strands ordered 32154, with β2 antiparallel to the other β-strands. Two of the three conserved helices are inserted between strands 3 and 4 and between strands 4 and 5 in the central sheet, while the third is typically near the C-terminus of the protein and positioned on the opposite side of the β-sheet. RNase H superfamily members are often characterized by four conserved acidic residues (the D,E/D,D,D motif) and a conserved histidine that populate the active site and support a 2-metal ion hydrolytic enzymatic mechanism (*26*). As described above, the CTD consists of a mixed 4-stranded β-sheet, with β2 running anti-parallel to β1, β3, and β4. In addition to losing a canonical strand in the central sheet, the CTD has only two α-helices, both at the C-terminus of the protein and both residing on only one side of the sheet. Further, several of the conserved catalytic/metal binding residues in the RNase H superfamily have been either been mutated to amino acids that do not fit the conservation pattern or reside on secondary structural elements that are not present in the CTD (Fig. 4E). Hence the CTD is likely a vestigial nuclease that has lost its hydrolytic capability.

Within the RNase H superfamily, the CTD is most structurally homologous to the nuclease domains from the large terminase proteins in other tailed bacteriophages (Fig. 4E). This result was somewhat unexpected since 1) the CTD was predicted to adopt an OB-fold; 2) the CTD has no recognizable sequence homology with the nuclease domain of non-phi29-like bacteriophages; and 3) phi29-like phages package a unit-length genome (capped at either end by a terminal protein; gp3 for phi29), and thus have no need for the nuclease activity. On the other hand, the result is not entirely surprising since all other non-phi29-like phages code for packaging ATPases where the CTD is a fully functional RNase H nuclease that cuts the concatenated genomic substrate at the beginning and end of packaging. One possible scenario is that the phi29 packaging motor evolved from other phage dsDNA packaging motors but lost the need for nuclease activity when the phage’s polymerase started making a unit length genome. As a result, selection pressure to conserve active site residues was removed and these amino acids could mutate without a fitness cost. Presumably, some secondary structural elements that positioned these residues also became expendable, allowing the phage to lose these structural elements as well.

### Characterizing CTD-nucleic acids interactions

#### Affinity of the CTD for DNA and pRNA

Despite the loss of nuclease activity, the observation that the phage maintained a minimal RNase H core suggests that some functional aspect of the RNase H fold has been conserved. Given the DNA translocating function of the motor and the need to assembly the functional ATPase ring on an RNA scaffold, conservation of a nucleic acid binding function seemed plausible. To test whether the CTD binds nucleic acids, we used fluorescence based binding assays to determine CTD binding affinities for dsDNA and a mimic of the pRNA A-helix. To quantify dsDNA binding, the fluorescence change of a fluorescein-labeled 22 base-pair dsDNA was monitored as a function of increasing protein concentration. The maximum observed value of fluorescence increased 30% by the end of the titration (Fig 5A). The resulting simple asymptotic curve is characteristic of a single binding site on the CTD for dsDNA and can be described Equation (1). A nonlinear least-squares fit to this equation yields a binding constant of K_1_ = 3.2 × 105 M^−1^ (K_d_ = 3.1 μM). Since the ATPase is known to bind to pRNA in the functional motor, we also sought to test whether the CTD interacts with the pRNA. It was not possible to concentrate pRNA to concentrations necessary for NMR experiments (described below). Therefore, to utilize a construct where we could directly compare fluorescence and NMR results, we designed a pRNA mimic that was could be chemically synthesized with a fluorescein tag. Thw construct is a hairpin structure that is predicted to folds into an RNA helix with the same sequence as the pRNA A-helix (Fig S4). Using a fluorescence assay similar to the one used for dsDNA, the fluorescein labeled pRNA mimic showed an ~30% rise in observed fluorescence intensity (Fig 5B) and indicated that the CTD binds the mimic with an affinity of K_1_ = 3.1 × 10^5^ M^−1^ (K_d_ = 3.2 μM). Given the similar binding affinities, these data indicate that isolated CTD does not have a binding preference for either dsDNA or the A-helix of pRNA. The relatively weak binding affinity for non-specific DNAs is not surprising given the translocating function of the packaging motor; binding must be strong enough to allow for interaction with DNA but not so strong that translocation would be stalled. Further, the full-length ATPase assembles as a pentamer on the pRNA via both the N-terminal domain and the CTD (Mao 2016 and below). Hence, the energies of five NTD and CTD binding events are summed for binding full-length gp16 to the pRNA pentamer, thus presumably even weak binding of the CTDs is sufficient in the context of the assembled complex of the full-length protein.

**Figure 5.**
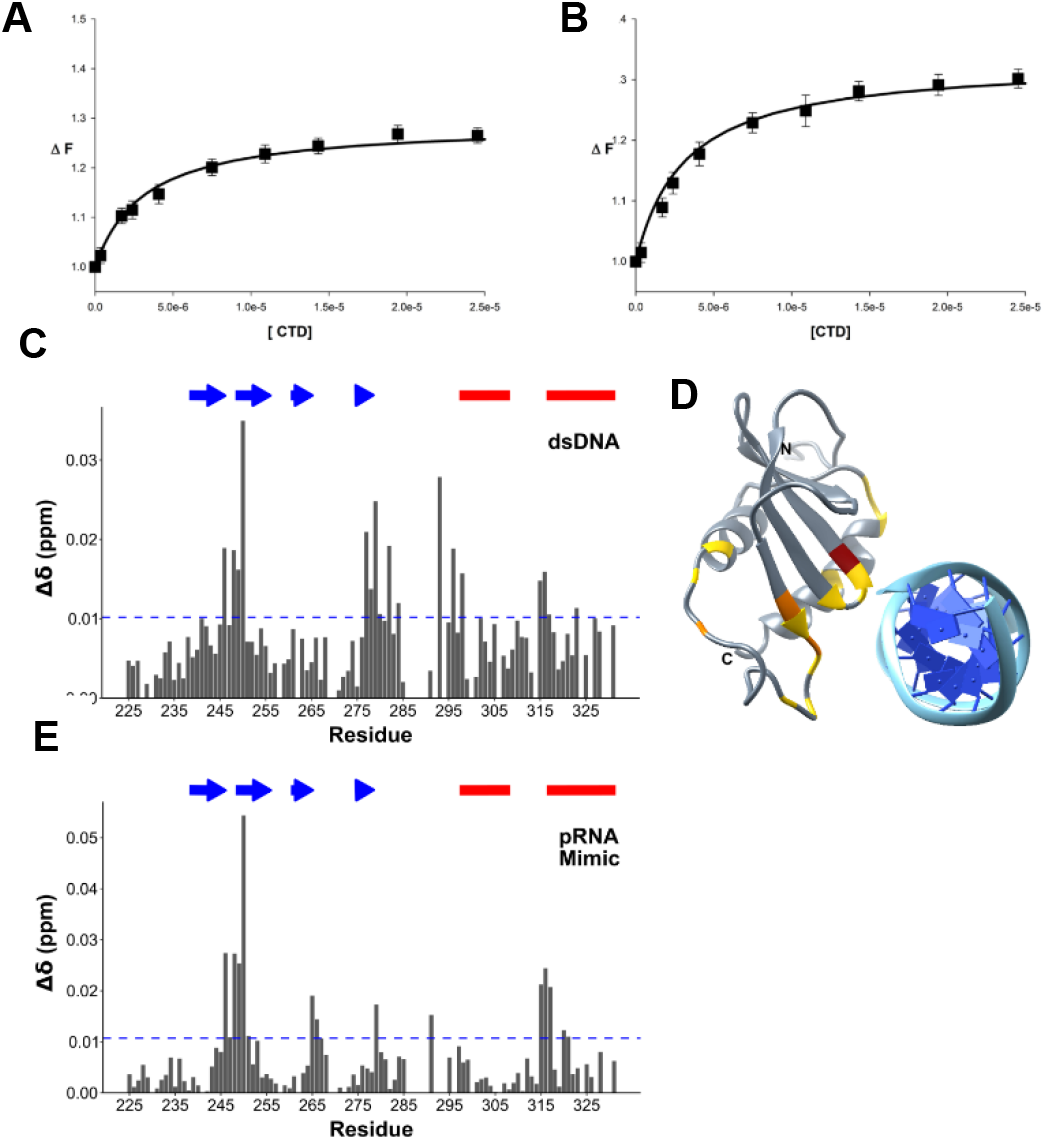
DNA and RNA binding to the CTD. (A) CTD interaction with dsDNA. (B) CTD interaction with the pRNA mimic. The solid lines are nonlinear least squares fits of the titration curves to equation 1 with binding affinities of Ka=3.2 × 10^5^ M^−1^, and K_a_=3.1 × 10^5^ M^−1^, for dsDNA and pRNA mimic respectively. (C) CSP profile of the CTD bound to dsDNA. The plot is shown with the significance cutoff of one standard deviation (blue dashed horizontal line) and the secondary structure elements, beta-strand (blue arrows) and alpha-helices (red bars) are highlighted above the plot. (D) CTD superimposed on RuvC bound to DNA substrate. The RuvC structure is removed from the figure for simplicity. The residues are colored according to standard deviations above null: 1 (yellow), 2 (orange) and 3 (dark red). (E) CSP profile of CTD bound to a pRNA A-helix mimic.

#### CTD residues that interact with DNA/pRNA

The binding constants of nucleic acids to CTD were too tight, (K_d_ < 10 μM) to determine accurately by NMR. However, specific residues in the CTD affected by nucleic acid binding could be identified by measuring their chemical shift perturbations (CSP) in HSQC spectra of 15N-labelled protein titrated with dsDNA, or the pRNA mimic. The results for dsDNA titration showed significant perturbations located in four regions: 1) between the β1 and β2 strands (residues 246-250); 2) at the end of β4 (residues 277-284); 3) before the beginning of α1 (residues 293-298); and 4) before the beginning of α2 (residues 315-317) (Fig 5C). The CSP profile using the pRNA mimic was strikingly similar the dsDNA-binding profile, except that the perturbations in residues around the beginning of the first helix were much smaller in magnitude (Fig 5D). While distant in sequence, the affected residues are close in structure space, mostly centered near the loop between the first two strands and the middle of α2.

To better understand how the CTD might bind nucleic acids, we superimposed the domain onto structures of human RNase H bound to an RNA:DNA hybrid (*27*)(PDB 2QKK) (Fig S5) and a nuclease related to large terminases, the *T. thermophiles* RuvC resolvase, bound to a Holiday junction (*28*)(PDB 4LD0) (Fig 5C,D). The alignments showed two different binding modes. The RNase H superimposition placed the CTD in the major groove of the hybrid with the edge of β4 contacting the 5’-3’ strand on one edge of the groove and α1 lying on the 3’-5’ strand at the other edge of the groove (Fig. S5). Based on this alignment, the C-terminus of our structure is positioned in the major groove. On the other hand, the RuvC alignment positioned the CTD over the major groove with the edge of the beta-sheet on the 5’-3’ strand, and the beginning of the second alpha-helix positioned over the complementary strand. Both superimpositions created a steric clash between the nucleotide substrate and the flexible loop joining β4 and α1; however clashing in the RNase H alignment is more extensive, as the entire loop is inserted between the two strands of the RNA:DNA hybrid. Further, the RuvC orientation is more consistent with the NMR CSP data, placing the loop residues with the largest perturbations (between β1 and β2) in direct contact with the DNA strand. The other residues with significant perturbations at the end of the last beta-strand and last helix are also in contact with the DNA. Hence, it seems likely that the binding interactions captured by NMR are more similar to the RuvC nucleic acid binding mode.

### Fitting NMR structure of the CTD into a cryoEM map of particles stalled during packaging

To understand how the CTD might interact with the pRNA and its dsDNA substrate during translocation, we fitted the NMR structure of the CTD into a cryoEM map of phi29 particles stalled during DNA packaging via the addition of the non hydrolyzable ATP substrate analog γ-S-ATP (**doi:** https://doi.org/10.1101/2020.05.23.112524). It should be noted that although the nominal resolution of the map was an ~5.2 Å, density corresponding to the CTD was considerably better. Visible side-chains and clear periodicity of a-helices suggest that the resolution for the CTD density is ~4-4.5 Å. In a previously published lower resolution reconstruction of particles stalled during packaging, five globular densities wedged between the translocating DNA and the pRNA were proposed to correspond to five copies of the CTD in assembled motor (*24*). This assignment is now confirmed by the unambiguous fit of our lowest energy NMR structure into corresponding density in the new higher resolution map (Fig 6A). In particular, the two α helices of the CTD fit well into clearly helical tubular densities in the map. Additionally, there is clear density connecting the N-terminus of our structure to density corresponding to the c-terminus of the fitted NTD. Further, density for the NTD-CTD linker region interacts extensively with neighboring ATPase subunits around the ring, consistent with our AUC experiments indicating that this region induces oligomerization in solution.

**Figure 6.**
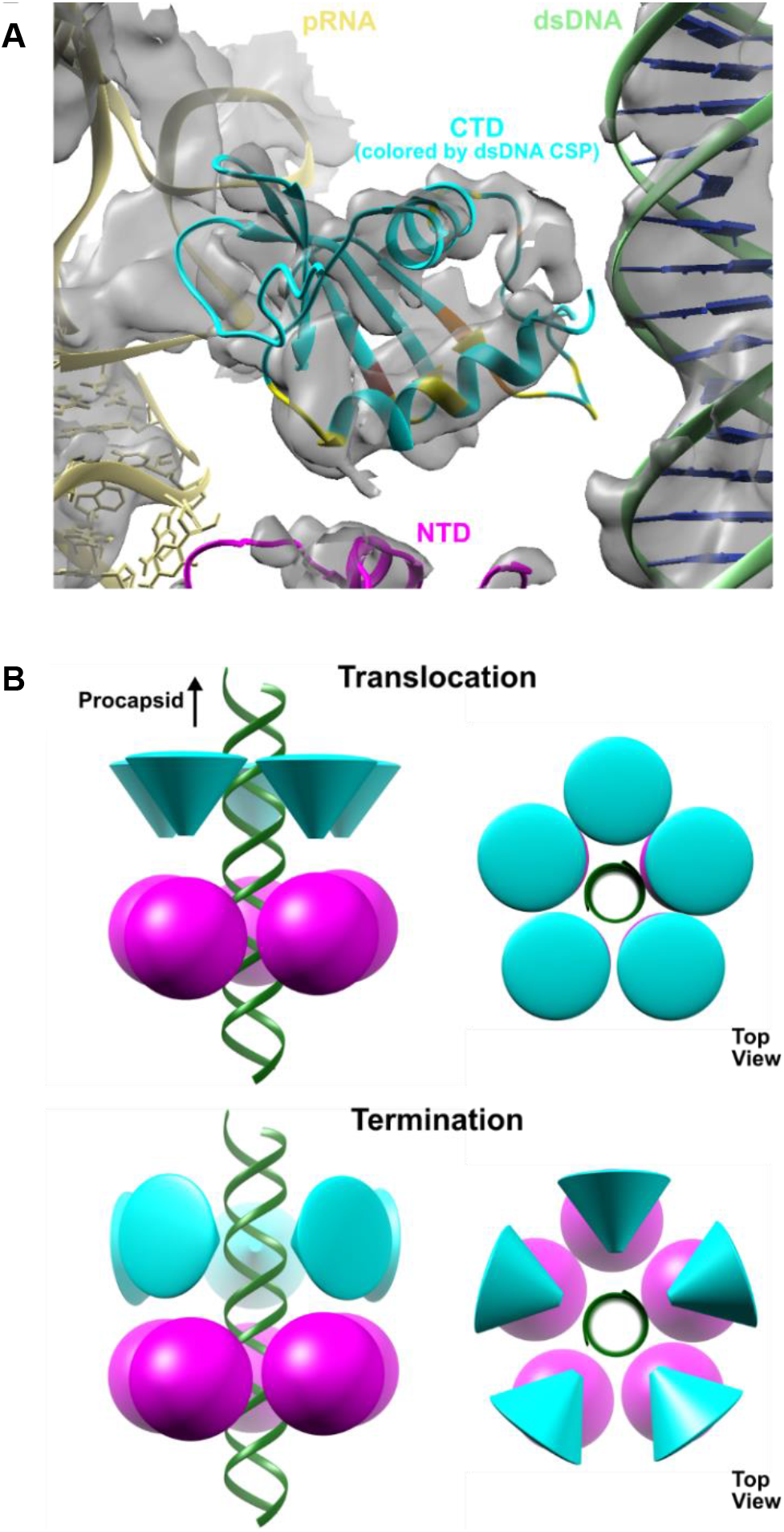
CTD fitted into a CryoEM Reconstruction of the phi29 Packaging Motor stalled during packaging. (A) The high resolution structures are fit into the density of a sub-nanometer cryoEM reconstruction of particles stalled during packaging (**doi:** https://doi.org/10.1101/2020.05.23.112524). The NMR structure of the CTD is shown in cyan with DNA-interacting residues identified in CSP data colored as in Fig 5D. The crystal structure of the NTD (PDB: 5HD9) is shown in magenta. The models of the pRNA (13) (PDB: 3R3F) and of B-form DNA are shown in tan and green respectively. (B) Schematic of the proposed transition from translocation to termination modes.

While the CTD is oriented with its N-terminus pointing towards the NTD, its C-terminus points towards the connector. CTD residues with side chains that are within 4 Å of the connector are located near the beginning of the first alpha helix (Y297, R300, K301). Residues potentially in contact with the pRNA are near the beginning of the first strand on the opposite face of the β-sheet (K236, K239, F240 and W254), and are conserved in phi29-like phages. The two helices of the CTD are oriented such that their ends point toward the DNA. For α1, this positions the positive end of the helix dipole close to the negatively charged DNA. This charge complementarity is further enhanced by side chain K294, which is close to the DNA. For helix α2, although the negative end of the helical dipole abuts the DNA, there are more positively charged residues, including R327, K328, and R330, resulting in a net positive charge. This fitting is consistent with packaging data for constructs where positively charged CTD residues close to the DNA were mutated; R327Q and R330Q decreased DNA packaging significantly, while K328N abrogated packaging entirely (data not shown). However, this orientation is seemingly contradictory to the NMR binding data, where the DNA seems to interact with the outer edge of the C-terminal helix (Fig 5D, F). This finding thus suggests that the NMR titration experiments capture a state of the motor that is structurally and functionally distinct from the state observed in cryoEM, and thus that the ATPase undergoes a substantial conformational change to rearrange into a translocation competent assembly.

## DISCUSSION

Packaging a viral dsDNA genome into a preformed protein capsid is a multi-step process requiring the action of a powerful molecular motor that can efficiently transition between packaging initiation, DNA translocation, and termination. To help clarify the roles of individual motor components during different stages of packaging, we determined the structure of the CTD of the phi29 packaging ATPase, probed its assembly and nucleic acid binding properties, and fit the structure into a cryoEM reconstruction of a functional motor complex assembled on procapsids. Despite the lack of any meaningful sequence homology, the CTD has a tertiary fold resembling an RNase H nuclease fold and is thus similar to the large terminase domains in other phages. In all other non-phi29 like phages, this domain is a fully functional nuclease that cleaves the dsDNA at the initiation and termination of packaging. However, presumably since phi29 packages a unit length genome, the CTD in phi29-like phages has lost this nucleolytic function (*4*). Despite the loss of nuclease activity, retention of a vestigial nuclease-like domain suggests that the CTD retains other functional properties common to both phi29-like and headful packaging phages.

### Initiation

Based on the data presented here, we propose that the different nucleic acid binding modes observed via cryoEM and NMR reflect two conserved yet distinct DNA binding functions; one necessary for translocation, and the other for termination/initiation. Further, we propose a model of the types of conformational changes that DNA packaging ATPase motors must undergo to transition between these two basic modes of operation (Fig 6B). The NMR CSP data shows that a nucleic acid binding interface resides near the loop connecting the first and second β-strands (Fig 5C,E). Superposition of the CTD onto the structure of the *T. thermophiles* RuvC resolvase bound to a Holiday junction suggests that this interface is similar to the ones used by other nucleases during strand cleavage, since the active site of the nuclease is near the scissile strand (Fig 5D,F). Presumably then, the nucleic acid binding interface identified via NMR and fluorescence corresponds to the initiation/termination interface.

### Translocation

In contrast, fitting the CTD into cryoEM density of packaging particles unambiguously shows an entirely different region of the CTD interacting with the incoming DNA. This interface is clearly a translocation interface. Further, in the cryoEM fit, the initiation/termination interface of the CTD is facing the NTD, occupying a position that assures it cannot interact with the dsDNA. In this orientation the CTD would have to rotate ~90 degrees to allow its termination interface to interact with the incoming nucleic acid (Fig 6). Thus, during translocation, the CTD termination interface is sequestered away from the DNA and positioned towards the NTD of the ATPase motor. A similar conformation in headful packaging phages would ensure that the genome is not prematurely cleaved during translocation of the genome into the procapsid.

Since the packaging reaction was stalled via the addition of γ-S-ATP, particles in this reconstruction presumably correspond to the end of the dwell; all five subunits have ATP bound (or the analog γ-S-ATP) and are poised to enter the burst. While the CTD is arranged as a planar ring in the reconstruction, the NTD adopts a helical configuration complementary to the dsDNA packaging substrate. The ATP-binding site is between residues such that trans-acting residues coordinate ATP binding, hydrolysis and product release. Based on this structure and other structures of phi29-like ATPases where the NTD is planar, we proposed that the motor cycles between helical and planar configurations of the NTD to package DNA (**doi:** https://doi.org/10.1101/2020.05.23.112524). During the burst phase, dsDNA is driven into the procapsid as the NTD helix collapses into a planar structure, driven by ATP hydrolysis and phosphate release. During the dwell, the NTD transitions back to a helical structure as ADP is exchanged for ATP sequentially around the ring. To ensure that DNA is not pulled out of the capsid during this transition, the NTD releases its grip on DNA. While this prevents the NTD from futilely moving the DNA back and forth as is cycles between helical and planar structures, it requires a mechanism to hold the DNA on place during the dwell as the motor resets. Hence, the CTD may function as a valve to prevent slippage of DNA during the dwell.

### Termination

After the motor protein receives the allosteric signal that the capsid is full, the CTD is rotated until the termination interface is positioned towards the DNA. In this position, the nuclease equivalent of the CTD in headful phages would not only be poised to cut the concatenated DNA genome, it would also be positioned to hold the DNA in place until the motor complex can dissociate from the filled capsid. Further, termination of one packaging event gives rise to the initiation complex of the next event in headful packaging phages. Therefore, this arrangement might also reflect the arrangement of motor components during initiation of packaging. Although phi29-like phages package a unit-length genome and thus wouldn’t move from procapsid to procapsid while attached to a freshly cut DNA, the termination interactions proposed here may also reflect initiation in phi29-like phages, providing a functional explanation for retention of a vestigial nuclease fold in the absence of any required nuclease activity.

A similar model for termination in the bacteriophage P74-26 packaging motor proposes that the nuclease activity of its terminase is sterically blocked by positioning the nuclease active site away from the incoming DNA during translocation, thus preventing premature cleavage. In this model the nuclease active site faces the portal vertex, and thus is not able to interact with the DNA during the translocation mode. After receiving a signal that packaging is complete, the nuclease domain is released from the portal vertex and free to bind and cleave the DNA (*29*). Other than the nuclease cleavage required in P74-26, the two mechanisms differ mainly in the orientation of the nuclease cutting interface prior to termination. In the structure presented here, the cutting interface during translocation is facing the NTD, whereas in the P74-26 model, the CTD is rotated ~180 degrees such that the cutting interfaces binds to the portal vertex. The differences between the proposed models may reflect different strategies of attaching the terminase to the procapsid; headful packaging phages presumably attach directly to the portal vertex, whereas phi29-like phages attach via the pRNA. While the relative orientations of the N- and C-terminal ATPase domains to the capsid has been the subject of some debate (*30*–*34*), the general conformational changes between initiation/termination and translocation modes described here would favor the arrangement where the CTD is closer to the portal vertex. Additional structural data of headful packaging motors attached to procapsids will further illuminate the motor conformations necessary for translocation and termination.

### Evolutionary relationship between CTD and pRNA

Despite a lack of sequence homology, structural homology between the CTD and the nuclease domains of other phage terminases suggests a shared evolutionary history. We suspect that the CTD in phi29-like phages evolved from nuclease domains in other phage terminases but lost nuclease activity concomitant with the need/ability to package a unit length genome. However, we cannot rule out the possibility that functional nuclease domains in other phage terminases evolved from a common nuclease-deficient ancestral phage similar to phi29-like phages. Regardless, the divergence of non-nucleolytic CTDs and functional terminase nuclease domains appears to be correlated to the presence or absence of pRNAs. Our fluorescence and NMR data suggest that the CTD does not discriminate between the A-helix of pRNA and dsDNA (Fig 5A, B). To prevent cleavage of the pRNA, the CTD must not have nuclease activity. Therefore, the loss of the nuclease active site may have facilitated the addition of the pRNA to the ATPase motor complex. Alternatively, as phi29-like phages moved to producing and packaging unit length genomes, the nucleolytic function of the CTD was no longer necessary. Hence, the CTD was able to evolve to the simplified vestigial nuclease fold we report here. However, a consequence of this loss of structural complexity may have been the introduction of new challenges regarding attaching the motor to procapsids. The C-terminal nuclease domains of other large terminases are believed to be involved in connecting the motor protein to the procapsid by binding to the portal vertex (*31*, *32*, *35*). The fitting of the CTD into cryoEM density of the motor vertex shows there are not substantial contacts between the CTD and the connector, and zero contacts with the portal-circumscribing capsid proteins. Instead, the pRNA is the major point of connection between the motor complex and the procapsid (*11*). Thus, one plausible scenario is that the loss of nuclease activity that allowed evolution of the simplified CTD also resulted in loss of the ability to bind the connector protein. In this scenario, the pRNA may have been recruited to reestablish a connection between the ATPase/terminase and the procapsid.

Alternatively, an evolutionary path wherein other phages once had but subsequently lost a pRNA-like molecule along with gain of nuclease activity cannot be ruled out. The RNA-world hypothesis proposes that on early earth, RNA served as both as the genetic material and as functional macromolecules in pre-biotic organisms (*36*). Thus, a primordial ancestor of other phages may have been similar to phi29-like phages but diverged evolutionarily. This divergence may have included the ability of large terminases to directly bind to the connector protein, thus, rendering any pRNA-like molecule unnecessary. Casting off the pRNA would have allowed the evolution of nuclease activity and replication and translocation of concatenated genomes seen in the majority of phages seen today. This evolutionary path implies that phi29 might more closely resemble a common ancestral phage. Indeed, phi29-like phages are the smallest known phages with the simplest genomes. Although it is not known when pRNAs first appeared in the virosphere, their recent detection in archaeal ampula viruses suggest their emergence predates the split between prokaryotes and archaea (*37*). While this path requires gain of two functions by an already present component (the CTD), the alternative path described above requires evolution of an entirely new component (pRNA molecule). Further experimentation and analysis will help resolve these and other questions.

## MATERIALS AND METHODS

### Protein Expression and Purification

The CTD of gp16 was cloned between the NdeI and XhoI sites of the pET30a plasmid. Protein was expressed in BL21(DE3) cells in either unlabeled LB or isotopically labeled M9 media. Purification was performed in 50 mM sodium phosphate at pH 7.5 for CTD-L (residues 208-332) and pH 6.5 for the truncated construct CTD (residues 223-332). The protein was initially purified using Talon resin (Clontech) with a linear gradient of 50 to 200 mM imidazole. The protein was further purified using a HiTrap Q HP column (GE Healthcare) for CTD-L and a HiTrap SP HP column for CTD with linear gradients of 50 to 500 mM NaCl. The final buffer was 20 mM sodium phosphate, pH 7.0, 50 mM NaCl for CTD-L. For CTD, the final buffer was 50 mM MES, pH 6.0. The buffer for ^13^C-labeled experiments contained 98% 13-^2^H MES in 100% ^2^H_2_O (Cambridge Isotope Laboratories, Tewksbury, MA).

### Analytical Ultracentrifugation

Sedimentation velocity experiments were performed in an Optima XL-A ultracentrifuge with an An-60 Ti rotor (Beckman Coulter, Indianapolis, IN) at 42,000 rpm and 30 °C to mimic the NMR experiments. The monitored wavelength was set maintain A ≈ 0.9 at each concentration. A partial specific volume of 0.73 and 0.74 mL/g was calculated by SEDNTERP for CTD-L and CTD, respectively (*38*). The buffer density and viscosity were also determined by SEDNTERP. Since MES did not exist in the SEDNTERP database, the default value for a generic buffer at 30 °C was used. The sedimentation velocity concentration profiles were modeled in SEDFIT to calculate the c(s) distribution (*39*).

### Fluorescence Experiments

Steady-state fluorescence titrations were performed using an ISS PC1 spectrofluorometer (ISS, Urbana, IL). Polarizers were placed in excitation and emission channels and set at 90° and 55° (magic angle), respectively, for all experiments in order to avoid possible artifacts due to fluorescence anisotropy of the sample. The nucleic acid binding of CTD was monitored by measuring changes in the signal originating from a fluorescein attached to the 3’ end of a strand in the dsDNA (GGTATCATGTGAGGGTGGGGGC-36FAM and GCCCCCACCCTCACATGATACC) or the 3’ end of the pRNA mimic (UCAAUGGUACGGUACUUCCAUUGUCAUGUUUUUCAUGGCAAAAGUGCACGCUACUUUGAU-36FAM) (λ_ex_ = 480 nm, λ_em_ = 520 nm). Binding curves were fit using KaleidaGraph software (Synergy Software, PA). Titration experiments were carried out by adding CTD to a solution containing labeled polynucleotide. The buffer, 50 mM sodium phosphate, pH 6.5, 200 mM NaCl, was used for all binding experiments. In these experiments the relative fluorescence change is defined as *ΔF*_*obs*_ = *Fi*/*F*_*0*_, where F_i_ is the fluorescence intensity at a given titration point and F_0_ is the initial value of the fluorescence of the sample. Both *F*_*0*_ and *F*_*i*_ were corrected for background fluorescence at the applied excitation wavelength (*40*).

The binding constants characterizing interactions between CTD and different nucleic acids substrates ([S] in equation) were determined using equation (1) below:

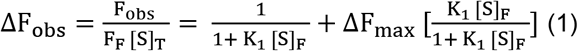

Where K_1_ is the binding constant characterizing the CTD association with substrates and ΔF_max_ is the maximum value of the observed relative fluorescence quenching.

### NMR Experiments

All data were collected at 303 K on a 600, 750 or 800 MHz spectrometer equipped with either a Bruker TCI or QCI-P CryoProbe (Bruker Biospin). All datasets were processed with NMRPipe (*41*) and analyzed with NMRViewJ (*42*). Backbone chemical shift assignments were determined from standard triple resonance experiments on ^15^N/^13^C-labeled CTD. Side chain assignments were obtained from ^15^N-TOCSY, HCCH-TOCSY and (H)CCH-TOCSY experiments. Distance restraints were determined from ^15^N-NOESY-HSQC (◻_mix_ = 120 ms) with ^15^N-labeled protein, and ^13^C-edited NOESY-HSQC (◻_mix_ = 120 ms) and 3D HSQC-NOESY-HSQC (◻_mix_ = 120 ms) on ^13^C-labeled CTD. Hydrogen bonding restraints were established from slow deuterium exchange. Backbone dynamics and rotational correlation times were determined by T1, T2 and heteronuclear NOE (hnNOE) experiments. Delays for the T1 experiment were 10, 80, 200, 330, 450, 600, 780, 1050, 1150, and 1250 ms, whereas those for the T2 experiment were 17, 34, 51, 68, 102, 119, 136, and 170 ms. The saturation delay for the hnNOE experiment was set to 3s. Rotational correlation times were obtained by fitting the average R_2_/R_1_ ratio of selected residues using an isotropic tensor in Tensor2 (refs). DNA and RNA titrations were performed with protein concentrations of ~100 μM in 50 mM sodium phosphate, pH 6.5, 200 mM NaCl, 0.02% NaN_3_, 2 mM Sodium 2,2-Dimethyl-2-silapentane-5-sulfonate (DSS). The dsDNA and pRNA sequences were the same as performed in the fluorescence experiments, except without the fluorescein label. All titration spectra were referenced to the proton chemical shift of the DSS methyl groups. The changes in chemical shift were calculated using equation (2):

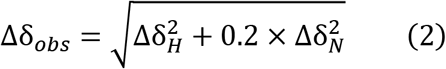

### Structure Calculation

The proton distance restraints were set with a lower limit of 1.8 Å and an upper limit of either 2.7, 3.5 or 5.0 Å based on NOE crosspeak intensities. Backbone phi and psi angles were predicted from chemical shift assignments using TALOS-N (*43*). The structure calculation was performed using a simulated annealing protocol implemented in XPLOR-NIH (*44*). Initially, 50 random structures were folded with torsion angle dynamics for 15 ps at 1000 K with van der Waals (VDW) force constant (0.002 kcal/Å^2^), VDW radii scaling (0.9), dihedral angles (200 kcal/rad^2^), NOE and hydrogen bond distance (0.50 kcal/Å^2^) restraints. A simulated annealing run from 1000 to 300 K was preformed, ramping the initial restraint values to 0.5 kcal/Å^2^ for the VDW force constant, 1.0 for the VDW radii scaling, 20.0 kcal/rad^2^ for the dihedral angles and 50.0 kcal/Å^2^ for both NOE and hydrogen bond restraints. After a 100 step torsion angle minimization, a refinement annealing step from 1000 to 20 K was performed while ramping the VDW force constant from 0.002 to 0.5 kcal/Å^2^, the VDW radii from 0.9 to 0.95, the dihedral angles from 200.0 to 20.0 kcal/rad^2^, NOE distances from 0.5 to 50.0 kcal/Å^2^, and hydrogen bonding restraints from 0.3 to 30.0 kcal/Å^2^. Angle and improper weights were ramped from 0.1 to 1.0 and 0.4 to 1.0 respectively for both annealing steps. The protocol ended with a 500-step torsion angle minimization and a 500 step Cartesian coordinate minimization. The NOE restraints were refined iteratively until the 15 lowest total energy structures had zero NOE violations. The quality of the structures was checked by Molprobity (*45*) and Chimera (*46*).

## ACKNOWLEDGEMENTS

This work is supported by NIH grants GM 122979 (to MCM and PJJ) and AI 087856 (to KHC). The authors also acknowledge the W. M. Keck Foundation and the John S. Dunn Research Foundation for funding the 800 MHz NMR spectrometer. Additionally, we are grateful for the helpful discussion and assistance provided by Krishna Rajarathnam, Junji Iwahara and Tianzhi Wang.

## Author contributions

BPM, PJB, HM and EAD performed the experiments. BPM and PJB performed the data analysis. BPM calculated the structure model. BPM, PJB and MCM wrote the manuscript. MCM and KHC supervised the research. PJJ helped interpret results and write the manuscript.

## NOTES

### Data and materials availability

The ensemble of structural models has been deposited in the PDB (6V1W) and the chemical shift data has been deposited in the BMRB (30691).

