## supplementary_materials for "NMR structure of a vestigial nuclease provides insight into the evolution of functional transitions in viral dsDNA packaging motors"

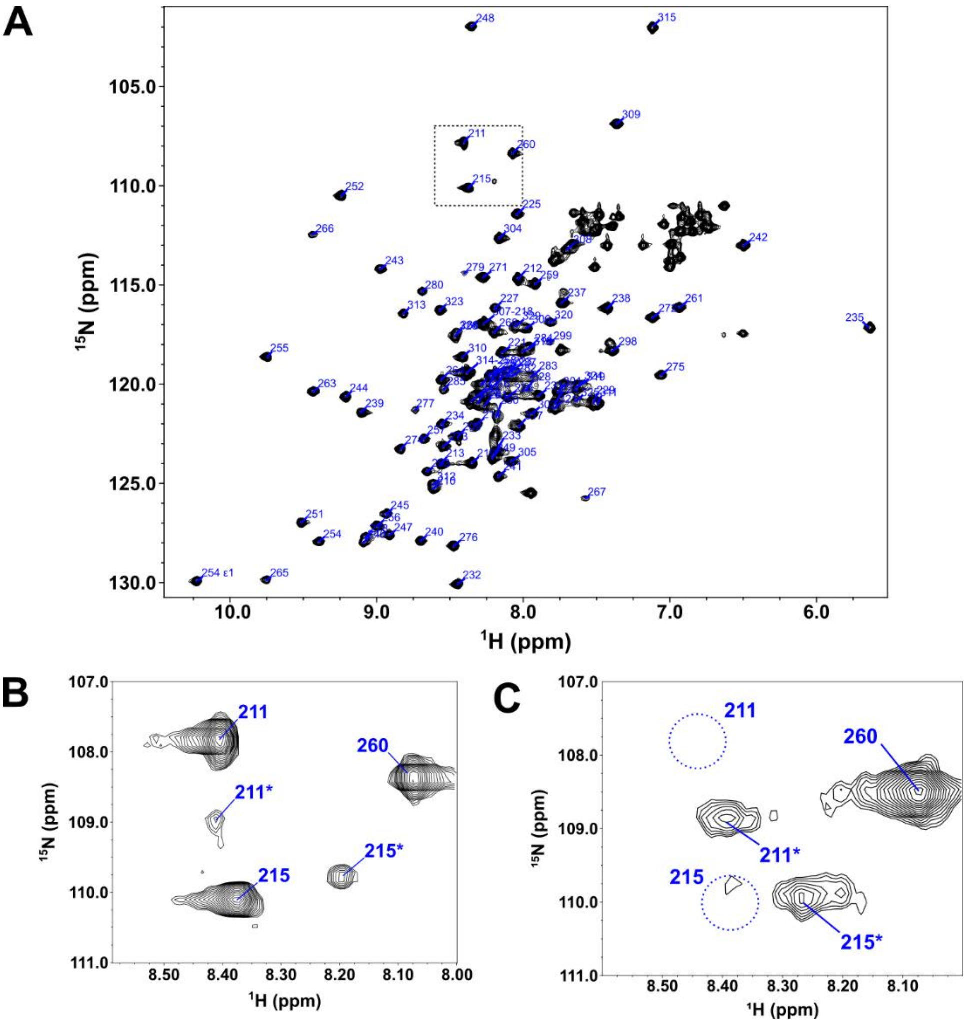


**Figure S1. NMR Spectrum of phi29 gp16 (208-332). (A)** The $^{1}$H-$^{15}$N HSQC of the phi29 gp16 (208-332) with backbone peaks labeled by residue number. The tryptophan side chain epsilon proton-nitrogen pair is also labeled. **(B)** Inset of the HSQC from the boxed region in (A). This region illustrates the extra peaks from the N-terminal tail described in the text. **(C)** An HSQC collected on an older gp16 sample. The region of the HSQC is the same shown in (B).

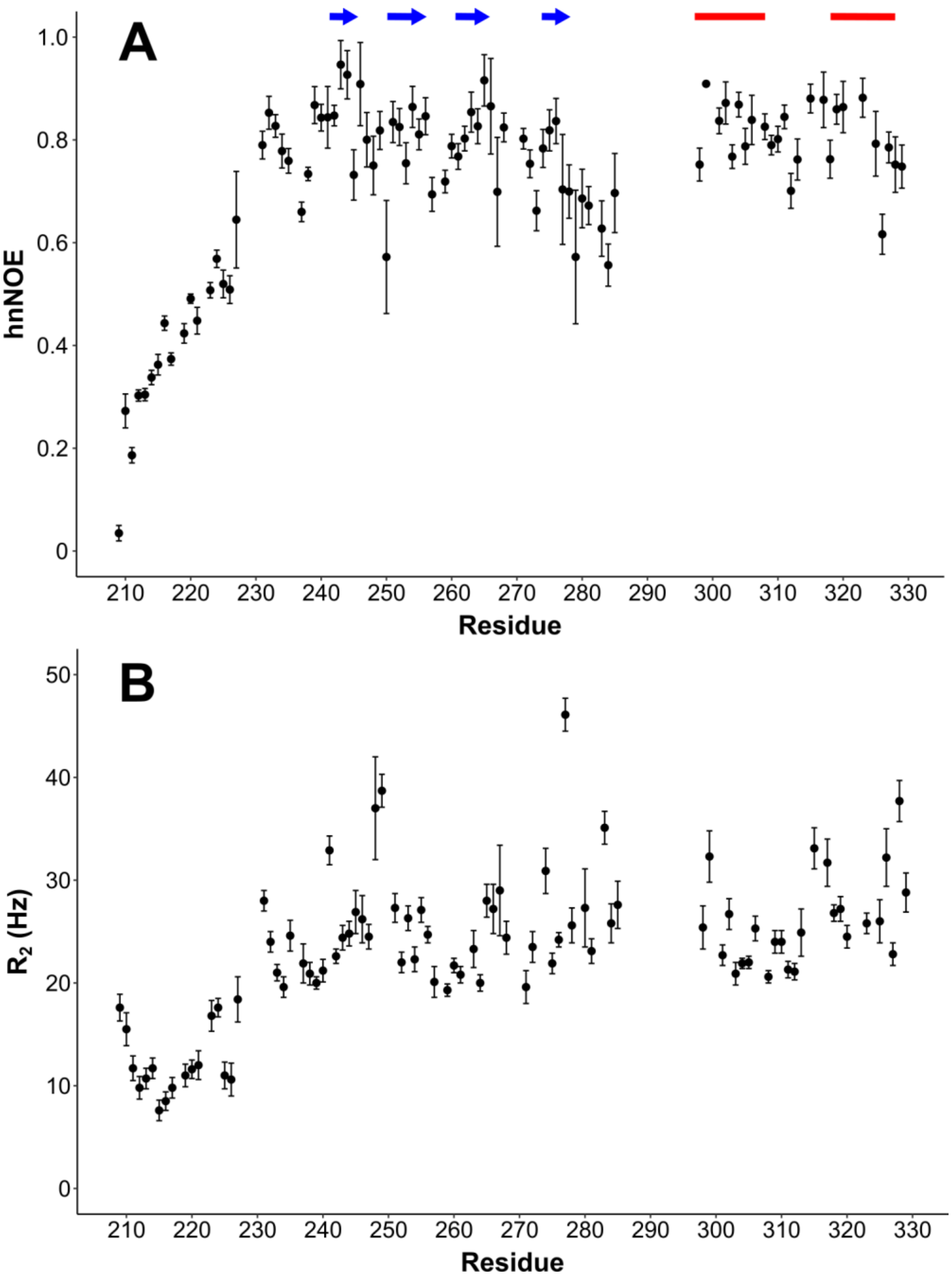


***Figure S2. Backbone Dynamics of phi29 gp16 (208-332).*** ***(A)*** *Heteronuclear NOE (hnNOE) of the CTD-L. This data is sensitive to fast time scale motions (ps-ns) with rigidity increasing with increasing hnNOE values. A hnNOE value $< 0.65$ is considered to be flexible.* ***(B)*** *Relaxation data from the CTD-L. Higher than average spin-spin relaxation rates ($R_2$) are indicative substantial signal broadening caused by exchanging structural states on NMR timescale (μs-ms). Residues in secondary structure elements are indicated by blue arrows (β-sheets) and red bars (α-helix) at the top of the figure.*

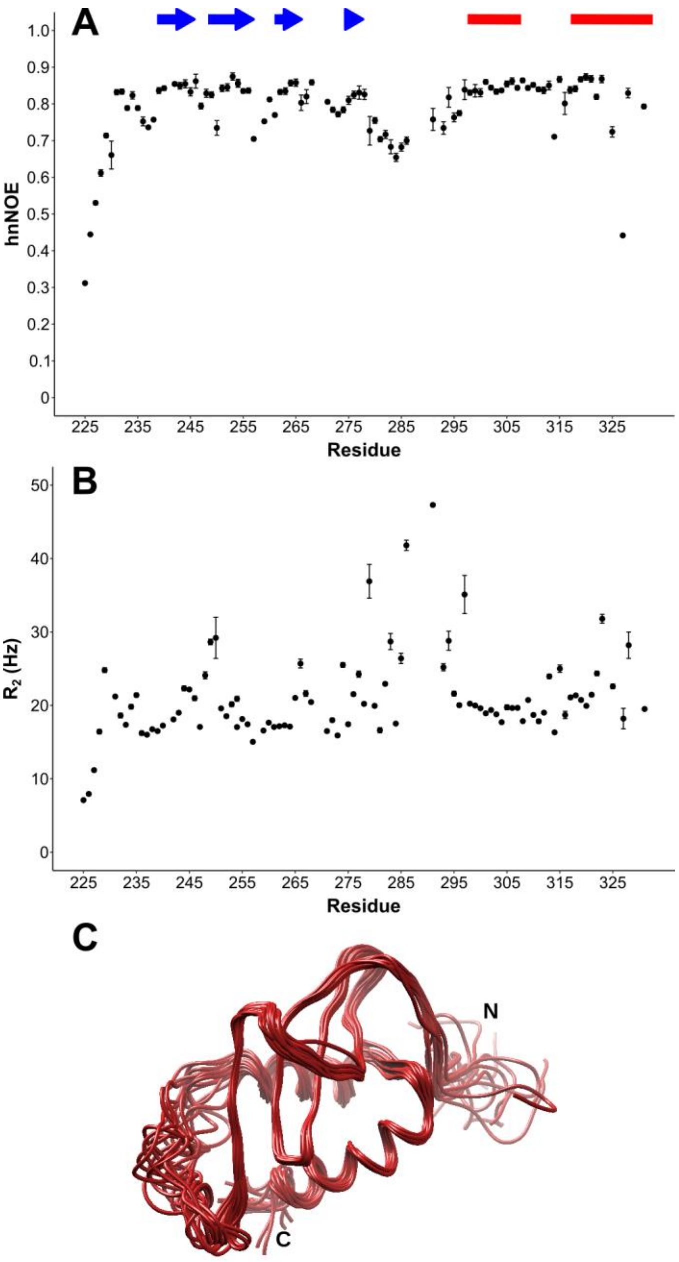


***Figure S3. Backbone Dynamics of phi29 gp16 (223-332).*** ***(A)*** *Heteronuclear NOE (hnNOE) of the CTD.* ***(B)*** *Relaxation data from the CTD. The data collectively highlight the flexibility of the N- and C-terminal tails, as well as the dynamics of the flexible loop between the 4$^{th}$ β-strand and the 1$^{st}$ α-helix (residues 282-296).* ***(C)*** *The 15-structure ensemble of the CTD.*

**Table S1 Structure Statistics for phi29 gp16 (223-332)**

| | |
|---|---|
| **Experimental Restraints** | |
| Long Range NOE (5 < i - j)[1] | 238 |
| Medium Range NOE (1 < i - j ≤ 5)[1] | 111 |
| Sequential NOE | 171 |
| Backbone Dihedral Angle | 212 |
| Hydrogen Bond | 44 |
| Total Restraints | 776 |
| **Ensemble Statistics (15 structures)** | |
| RMSD Distance and Ideal Geometry Restraints | |
| Distance (Å) | 0.061±0.002 |
| Bond Length (Å) | 0.016 |
| Bond Angle (°) | 0.967±0.017 |
| Bond Improper (°) | 0.508±0.029 |
| Ramachandran Statistics (MolProbity) | |
| Favored Region (%) | 91.4 |
| Allowed Region (%) | 98.5 |
| Disallowed Region (%) | 1.5 |
| RMSD from Average Structure | |
| Backbone Atoms (Secondary Structure)[2] | 0.58 |
| All Heavy Atoms (Secondary Structure)[2] | 1.1 |

[1]i-j is the distance of two residues in sequence space

[2]Secondary structures residues are defined as 242-245,250-256,261-265,274-278,297-308,318-328

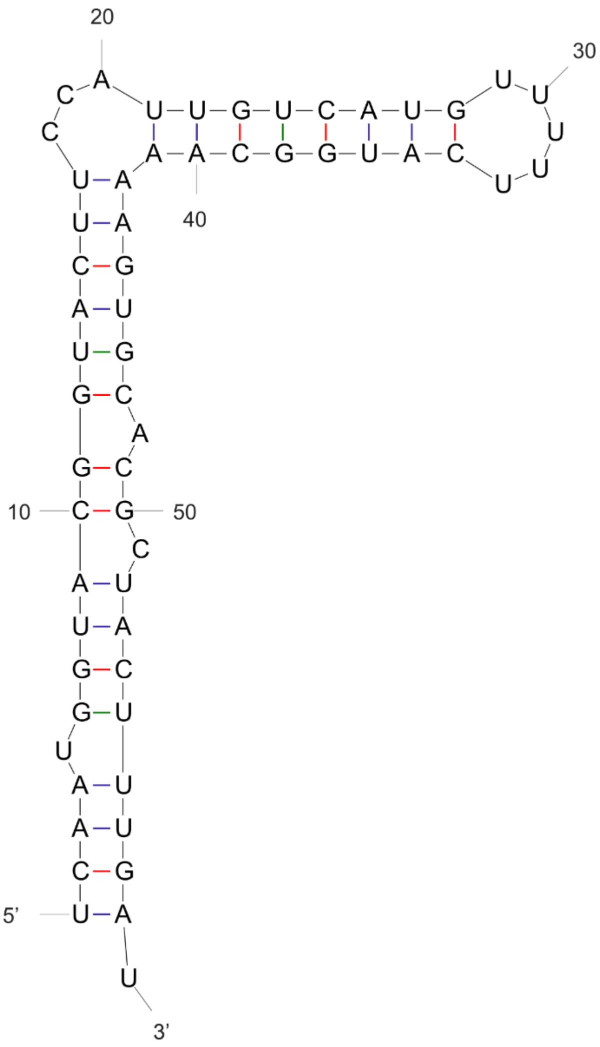


*Figure S4. A secondary structure diagram of the pRNA mimic. The secondary structure was predicted using the mFold web server (1).*

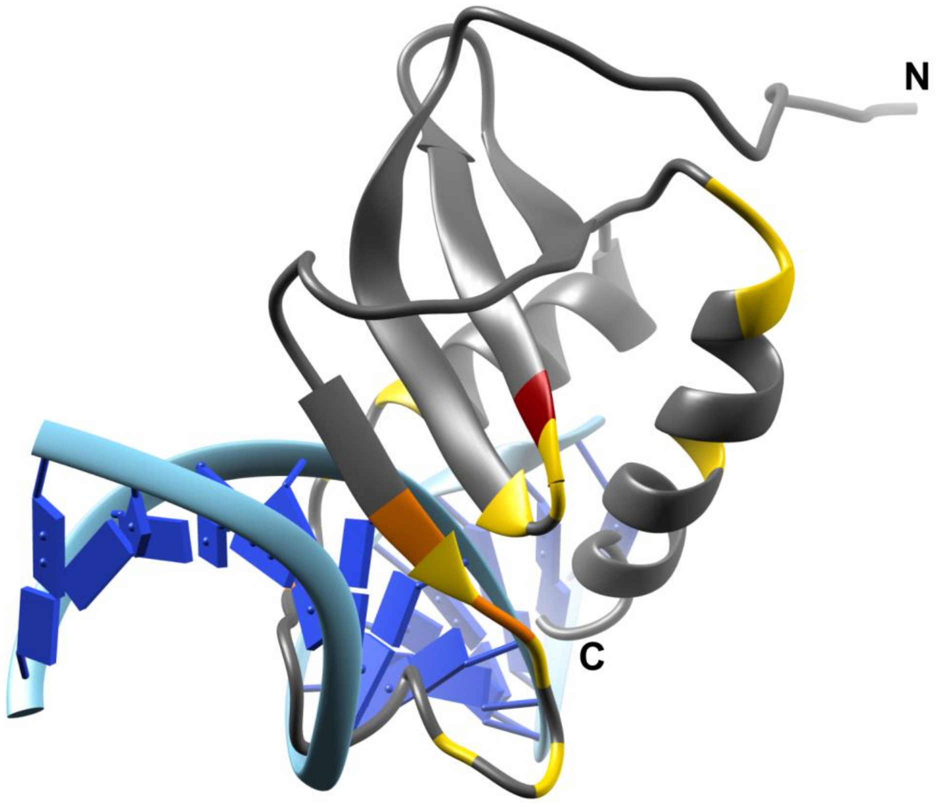


***Figure S5. Superpostion of CTD over RNase H bound to its Nucleic Acid Substrate (PDB: 2QKK).***

*The structure of the RNase H protein was removed after superimposing the CTD on it for simplicity. The residues are colored by their CSP with dsDNA bound (Fig. 5C): >1 std. dev. (yellow), >2 std. dev. (orange), >3 std. dev. (red).*

### Supplemental Material References

1. M. Zuker, Mfold web server for nucleic acid folding and hybridization prediction. *Nucleic Acids Res.* **31**, 3406-3415 (2003)
